# Connectivity analysis of GEF/GTPase networks in living cells

**DOI:** 10.1101/728998

**Authors:** D.J. Marston, M. Vilela, Jinqi Ren, George Glekas, Mihai Azotei, G. Danuser, J. Sondek, K.M. Hahn

**Affiliations:** Dept of Pharmacology, UNC Chapel Hill, Chapel Hill, NC; Departments of Bioinformatics and Cell Biology, University of Texas Southwestern Medical Center, Dallas, TX.

## Abstract

The cytoskeleton is regulated by dynamic, multi-layered signaling networks that interconnect Rho family small GTPases with exquisite spatiotemporal precision^1^. Understanding the organization of these networks is challenging, as protein activation and interaction occur transiently and with precise subcellular localization. We and others have used fluorescent biosensors in living cells to map the activation patterns of the Rho family small GTPases relative to the changes in cell edge dynamics that they produce^2–4^. These GTPases are controlled by the localized activity of numerous Rho guanine nucleotide exchange factors (RhoGEFs) with overlapping GTPase specificity^5,6^. Here we extend this analysis to determine functional relationships between GEFs and GTPases in controlling cell edge movements. First, biosensors for GEF activation are produced, and their activity correlated with edge dynamics. We then shift the wavelengths of GTPase biosensors to image and correlate the activation of GEFs and GTPases in the same cell. Using partial correlation analysis, we can parse out from such multiplexed data the contribution of each GEF – GTPase interaction to edge dynamics, i.e. we identify when and where specific GEF activation events regulate specific downstream GTPases to affect motility. We describe biosensors for eight Dbl family RhoGEFs based on a broadly applicable new design strategy (Asef, Tiam1, Vav isoforms, Tim, LARG, and β-Pix), and red shifted biosensors for RhoA, Rac1 and Cdc42. In the context of motility, functional interactions were identified for Asef regulation of Cdc42 and Rac1. This approach exemplifies a powerful means to elucidate the real-time connectivity of signal transduction networks.

---

In previous studies of Rho GTPases in motility we could determine the relative timing and location of Rac1, Cdc42, RhoA, and RhoC activation by first using fluorescent biosensors to correlate the activation of each GTPase with cell edge movements, and then using cell edge movement as a common fiduciary to relate the different GTPases to one another^3,7,8^. This allowed us to predict the relative spatio-temporal dynamics of the four signaling activities with exquisite precision, but we could not predict whether any of the signals were directly or indirectly coupled, how specific source signals contributed to the total modulation of each target signal, and the effects of specific couplings on downstream events. Answering such questions is essential to further understand the complex relationships between GEFs and Rho GTPases: One GTPase may be activated by multiple GEFs, even in the same location, and one GEF may activate multiple GTPases. Thus, statistical approaches are required to quantify the fraction of a GTPase signal that results from a particular GEF, and the relative contribution a GEF makes to each of the downstream GTPases it interacts with. Moreover, to appreciate the functional diversity of a GEF in controlling, e.g., cell motility vs other downstream functions, it is necessary to statistically determine the fraction of cell edge dynamics that results from a particular GEF-GTPase interaction.

To perform these statistical analyses, it was first necessary to produce biosensors that report the activity of GEFs in living cells. Many Dbl family RhoGEFs are regulated through occlusion of the GTPase binding interface by an autoinhibitory domain (AID)^5^, providing a route to Dbl family biosensors. In our first biosensor target, the RhoGEF Asef, an SH3 domain acts as the AID; it undergoes a conformational rearrangement when adenomatous polyposis coli (APC) binds to the upstream ABR (APC binding region), leading to Asef activation^9^ (Figure 1a). We produced an Asef analog that reports these activating conformational changes by inserting two fluorescent proteins into the flexible linker between the SH3 AID and the catalytic DH domain. The activating conformational changes affected FRET between the fluorescent proteins by altering their distance or relative orientation. We used high-throughput microscopy^10^ to test insertion of Cerulean^11^ and Venus^12^ fluorescent proteins at a series of positions between the AID and DH domain, optimizing FRET intensity and the activation-dependent difference in the donor/FRET emission ratio (**Extended Data Figure 1a).** We then screened a small library combining different fluorophore pairs (Cerulean3^13^, TagCFP^14^, or mTFP^15^; combined withYPet^16^ or a series of YPet circular permutations) (**Extended Data Figure 1a**). This led to a biosensor with 75% difference in donor/FRET ratio for wild type versus constitutively active Asef (Figure 1a). Similar changes were seen upon co-expression of known activating proteins, a fragment of APC^9^ or constitutively active Src^17^ (Figure 1b). No ratio change was observed when kinase-dead Src or a non-binding APC mutant were used, or when the FRET pair was moved to a site in the RhoGEF that does not undergo a conformational change (Figure 1b).

**Figure 1.**
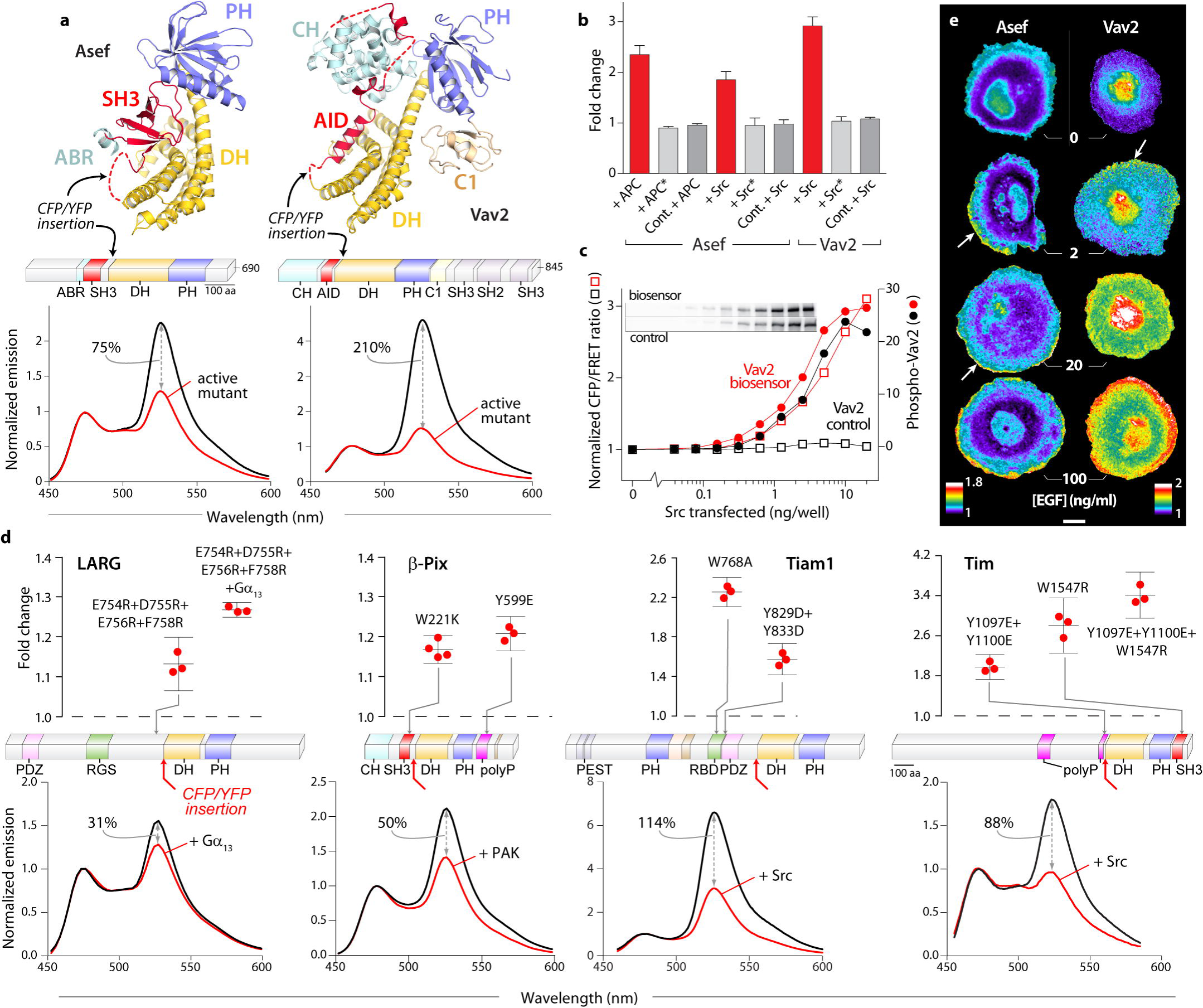
**(a)** Biosensors for Asef and Vav2. Autoinhibited structures of Asef and Vav1 (PDB entries 2PZ1 and 3KY9, respectively, top) and domain architecture of full length Asef and Vav2 (ABR - APC binding region, AID - autoinhibitory domain, C1 - C1 domain, CH - calponin homology, DH - Dbl homology, PH - pleckstrin homology, SH2 - Src homology 2, SH3 - Src homology 3, middle) with insertion sites of FRET pair cassettes indicated. Emission spectra (λ_ex_ = 430 nm) of wild type and active forms (Asef - V252E, Vav2 - Y142E:Y159E:Y172E) expressed in HEK293t cells (bottom). Change in donor/FRET ratio upon activation is indicated. **(b)** Stimulation of Vav2 and Asef biosensors using Src and APC in HEK293t cells. Src* is kinase dead, APC* does not bind to Asef. In biosensor control (Cont.) the FRET pair is moved to the C-terminus. Ratio normalized to empty vector control. **(c)** Effect of increasing Src levels on donor/FRET emission ratio (open squares) and phospho-Tyr^172^ levels (solid circles) of Vav2 biosensor (red) and control (black). Normalized to empty vector transfection control. Inset – phospho-Vav blots. **(d)** Biosensors for LARG, β-Pix, Tiam1 and Tim. Domain structures show site of FRET pair insertion in full length RhoGEFs (PDZ – PSD95/Dlg1/zo-1; RGS – regulator of G protein signaling; polyP – polyproline; PEST – Sequence rich in amino acids P, E, S and T; RBD – Ras-binding domain; other domains are as listed above, middle). Lower graphs show emission spectra of wild type biosensors +/− indicated activators (λ_ex_ = 430 nm), change in donor/FRET ratio upon activation is indicated. Upper graphs show ratio change caused by indicated mutations as compared to wild type. **(e)** A431 cells expressing Asef (left) and Vav2 (right) biosensors after stimulation with increasing amounts of EGF. Arrows point to high ratio values at the edge of cell protrusions. Ratios in pseudocolor normalized so lowest value = 1. Scale bar 10µm.

We hypothesized that this approach could be applied to other RhoGEFs that undergo a conformational change upon release of autoinhibition, so we tested Vav2, where the DH domain is blocked by an upstream helical AID^18^ (Figure 1a). Autoinhibition is relieved when Src and other kinases phosphorylate tyrosines in the AID^19,20^. As with Asef, we optimized the site of fluorophore insertion between the AID and DH domains, and screened fluorophore combinations (**Extended Data Figure 1a)**. In addition, we tested fluorophore-fluorophore connectors of varying length and rigidity to impose constraints on the conformation of the inserted segment (**Extended Data Figure 1b**). This led to three biosensors, whose fluorescence ratio changed 210% (TagCFP donor), 185% (mCerulean3 donor), and 105% (mTFP donor) upon activation (Figure 1a, **Extended Data Figure 1c**). The detectability of GEF activity was a function of both the extent of fluorescence change and the brightness of the fluorophores. In the imaging studies below, we used the brightest donor (mTFP) even though it produced less change (**Extended Data Figure 1c**).

To test whether the Vav2 biosensor could report activating conformational changes in living cells, we compared it and the non-responsive control biosensor in HEK293 cells, examining response to increasing amounts of constitutively active Src. Dose-dependent phosphorylation of the biosensor and the control biosensor were equivalent, as shown by blotting with a phospho-Tyr174 antibody, but only the real biosensor showed an increase in fluorescence emission ratio (Figure 1c). As with Asef, this ratio change was not seen with inactive Src (Figure 1b). Using the optimized Vav2 biosensor as a template we also produced biosensors for other Vav family members simply through limited screening of the insertion site (**Extended Data Figure 1d**).

For Asef and Vav2, high-resolution crystal structures were available to identify AID interactions. We next attempted to make biosensors for RhoGEFs proposed to have autoinhibitory regulation, but where structural information was limited. The RhoGEF Tim contains a putative helical region that is thought to directly interact with the DH domain^21^, equivalent to the Vav2 AID, and autoinhibition is maintained by polyproline and SH3 domains that flank the DH domain^22^: For the GEFs Tiam1 (T-cell lymphoma invasion and metastasis 1) and LARG (Leukemia-associated RhoGEF), Small-angle X-ray scattering suggests that an N-terminal domain folds over the DH domain^23,24^, and binding sites for known regulators lie upstream of the DH domain (Ras^25^/tyrosine kinases^26–28^ for Tiam1, and Gα_13_^29^ for LARG). The RhoGEF β-Pix has a polyproline-SH3 domain pair flanking the DH domain and multiple sites downstream that are proposed to regulate activity^30^. For each of these RhoGEFs we screened donor/acceptor insertion sites directly upstream of the DH domain, and optimized the response as before (**Extended Data Figure 1a**). The final set of biosensors had dynamic ranges varying from 31% for LARG, to 115% for Tiam1 (Figure 1d). For Tim, mutations within the autoinhibitory helix that mimic Src phosphorylation, and mutations within the SH3 domain that prevent polyproline binding, caused changes in the donor/FRET ratio as large as those produced by Src co-expression. The combined mutations had an additive effect (Figure 1d).

The new LARG, Tiam1 and β-Pix biosensors provided insight into potential regulatory mechanisms. Specific residues within the RBD (Ras binding domain) of Tiam1 caused large FRET changes, suggesting that this domain has an autoinhibitory role (Figure 1d). Mutations of negatively charged residues in LARG^31^ and phosphorylation sites in β-Pix that were thought to be involved in RhoGEF regulation induced modest FRET changes, and mutation of the SH3 domain in β-Pix also affected FRET, suggesting that there may be an autoinhibitory role for the polyproline-SH3 domain pair flanking the DH domain, as there is for Tim (Figure 1d).

We validated the new biosensors by examining their response to known stimuli in living cells. In response to EGF, Vav2 and Tiam1 showed widespread activation (Figure 1e, **Extended Data Figure 2a**) while Asef was activated only within 2µm of the cell edge (Figure 1e). LARG and β-Pix showed reversible activation concentrated at the leading edge of randomly migrating cells (**Extended Data Figure 2c**). More detailed studies of Vav2 showed that its response was dose dependent and reversible, with activation within 30 seconds (**Extended Data Figure 2b**).

**Figure 2.**
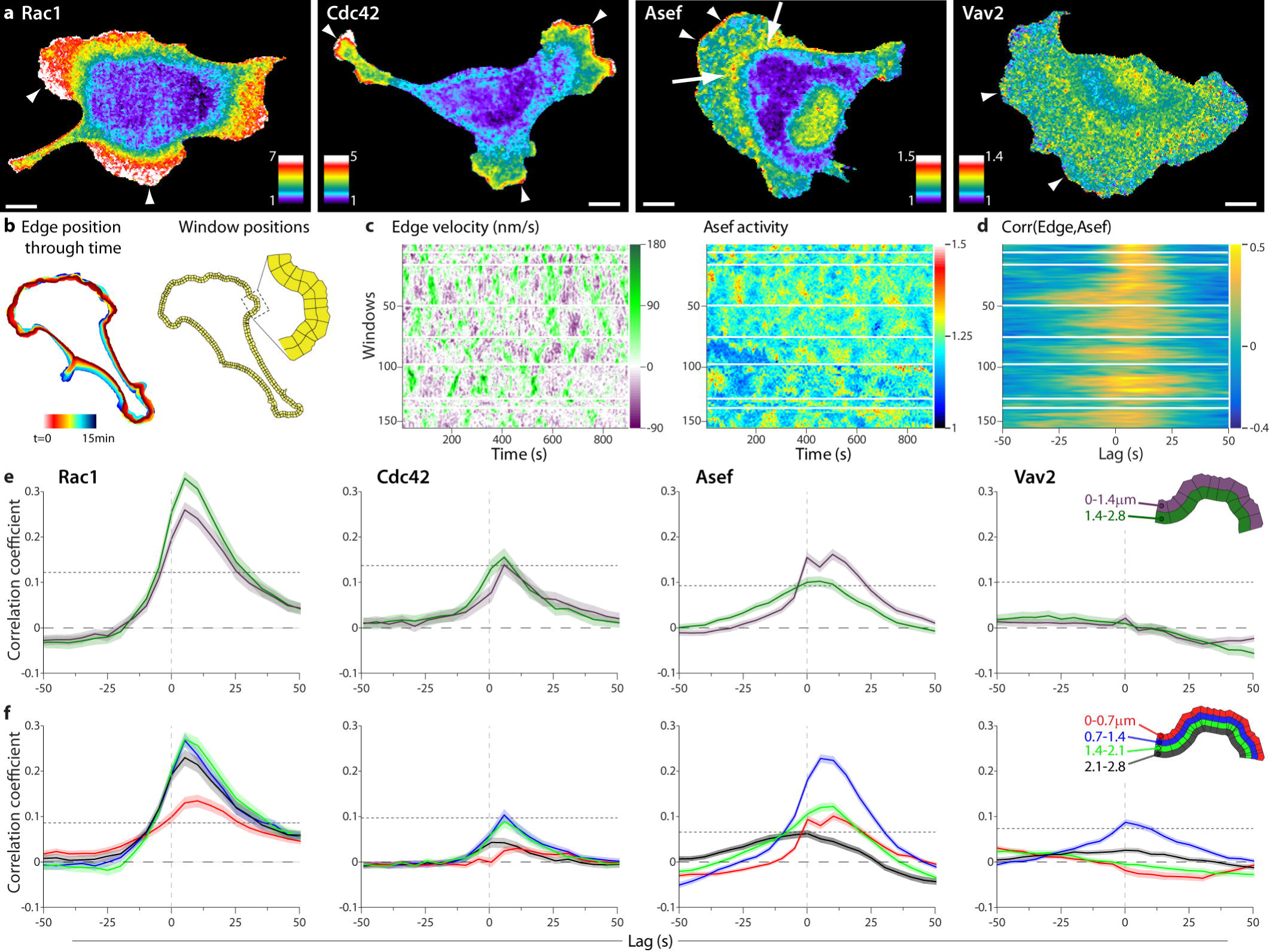
**(a)** Rac1, Cdc42, Asef, and Vav2 activation reported by biosensors in MDA-MB-231 cells undergoing random edge motion. Pseudocolor as in Fig. 1. Scale bars 10µm. **(b)** Evolution of cell edge positions (color-encoded from red (early) to blue (late) time points, left). Two rows of sampling windows, each 1.4 μm deep placed at the cell edge. **(c)** Maps of edge velocity (left) and Asef biosensor activities (right) along the edge. Green regions are protruding, purple regions retracting. Red/yellow regions have high activity, blue regions have low activity. Each column is a single time point. White horizontal bars show quiescent regions that are excluded. Scale bars are shown to right of each map (X units of velocity, X units of activity). **(d)** Cross correlation coefficients between edge velocity and Asef activity. Gold shows high correlation and blue negative correlation. Each row is a single position along the edge corresponding to the activity maps. **(e)** Average cross-correlation functions for each biosensor. Analysis using 1.4 µm window size. (n cells, m windows); Rac1 (9, 259); Cdc42 (6, 204); Asef (8, 448); Vav2 (6, 360). **(f)** Average cross-correlation functions for each biosensor. Analysis using 0.7 µm window size. (n cells, m windows); Rac1 (9, 518); Cdc42 (6, 408); Asef (8, 896); Vav2 (6, 720). Inset shows window size and color key.

For studies of GEF-GTPase circuitry, we focused on GEFs that were stimulated by EGF, to examine protrusion/retraction cycles during EGF-induced chemokinesis. Biosensors for Asef, Vav2, and improved versions of our previously published biosensors for Rac1^3^ and Cdc42^32,33^ (**Supplemental methods and Extended Data Figure 3a**) were stably expressed in MDA-MB-231 cells and studied in EGF-containing medium. Biosensors were kept below expression levels that perturbed motility behaviors (**Extended Data Figure 4**). As previously described, Rac1 (Figure 2a, **Supplemental Movie 1**) and Cdc42 (Figure 2a, **Supplemental Movie, 1**) showed broad gradients of activity dropping from the cell edge to 5 – 8 µm behind the edge, consistent with activation in protruding lamella (**Extended Data Figure 5a, b**). The new biosensor showed that Asef activation was generally restricted to a narrow band at the leading edge of protrusions (resembling that produced by acute EGF stimulation of A431 cells, Figure 1e), together with a broader region 5-10 microns back from the edge, at the base of lamella, (Figure 2a, **Extended Data Figure 5a, b Supplemental Movie 1**). The latter activation colocalized with Rab-7-mScarlet, suggesting activated Asef interacts with a sorting compartment (**Extended Data Figure 5c**). When we moved the FRET pair in the Asef biosensor to a region that should not undergo conformational change, activation at both cellular locations was eliminated (**Extended Data Figure 5d**). In contrast, Vav2 showed diffuse activation throughout the cell, with only a few ‘hot spots’ adjacent to the edge (Figure 2a, **Supplemental Movie 1**).

**Figure 3.**
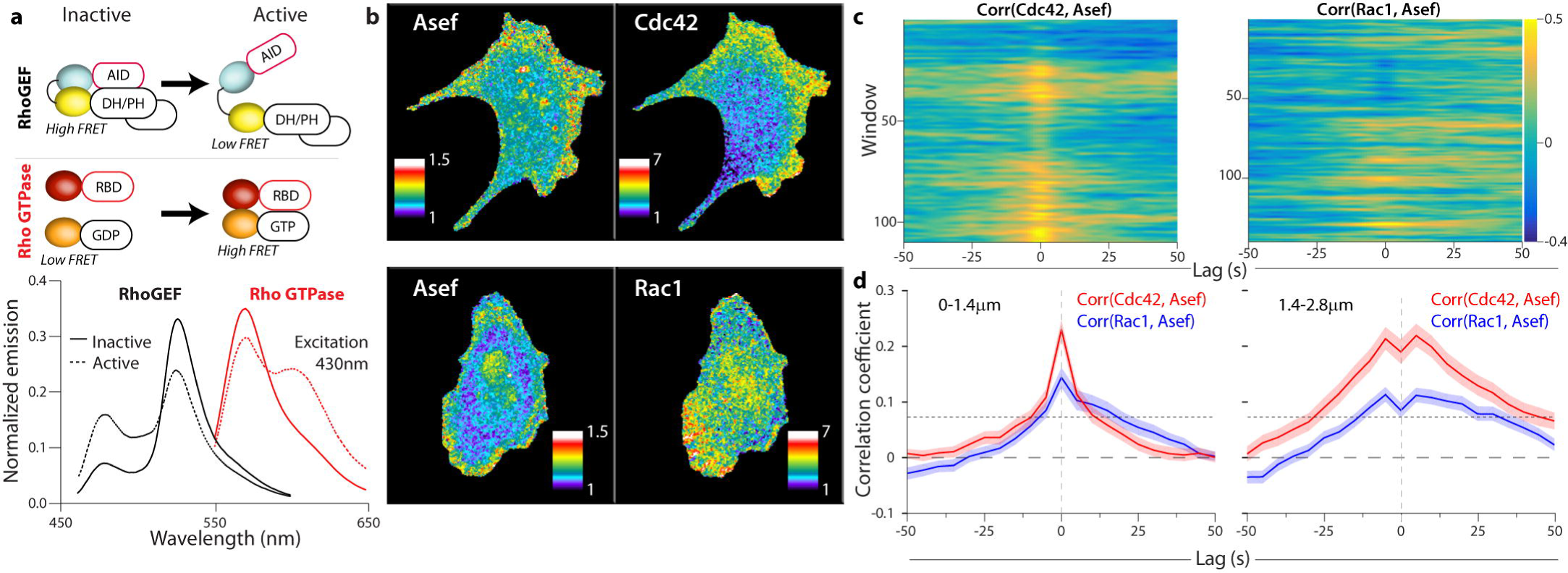
**(a)** Biosensor designs and fluorescent proteins used for multiplexing (upper). Emission spectra of representative RhoGEF and Rho GTPase biosensors (lower). Inactive biosensors in solid lines, activated forms in dotted line. The two spectra were obtained using the same excitation wavelength (430 nm) **(b)** RhoGEF and Rho GTPase biosensors imaged simultaneously in MDA-MB-231 cells undergoing constitutive edge motion. **(c)** Cross correlation coefficients between Rho GTPase activity and Asef activity in the 1.4-2.8 µm layer. Gold shows high correlation, blue negative correlation. **(d)** Average cross-correlation functions for each biosensor combination; (n cells, m windows); Cdc42 (5, 729) (left); Rac1, (6, 719) (right)

**Figure 4.**
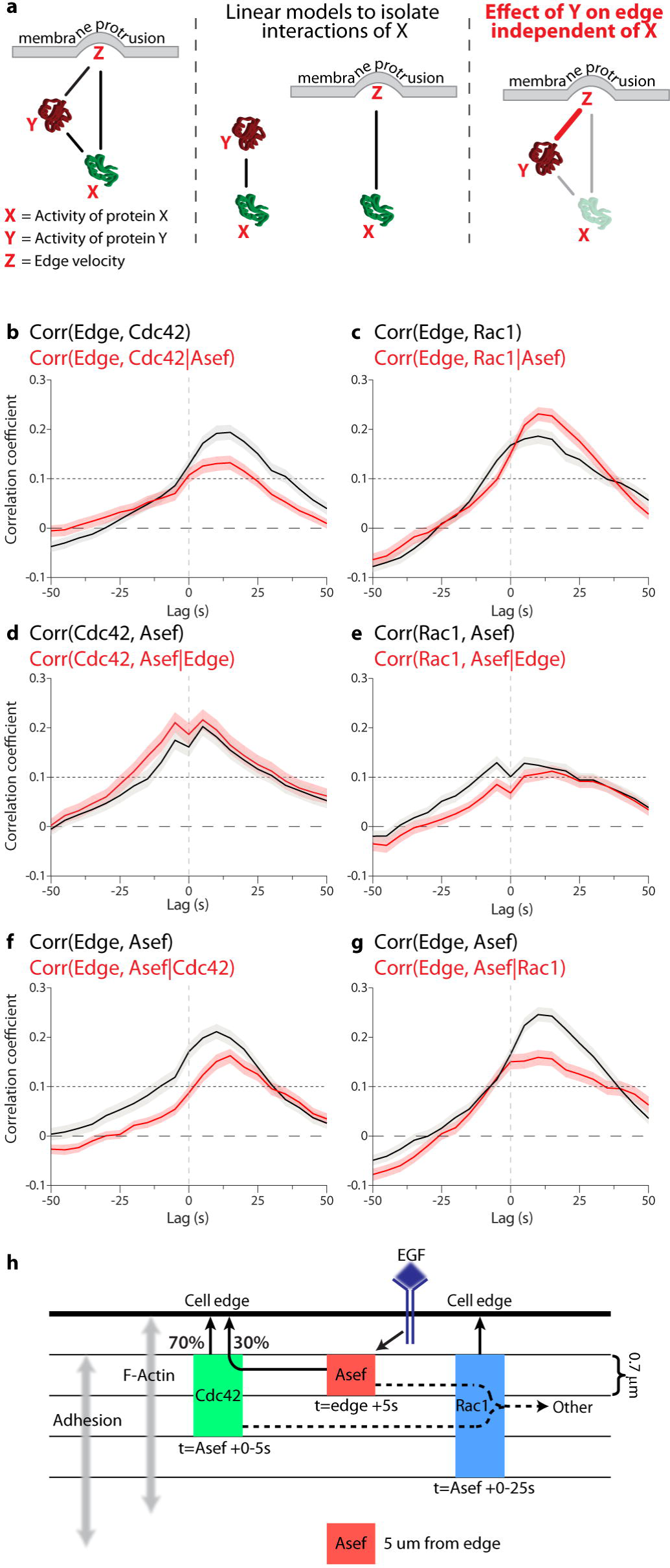
**(a)** Model for partial correlation analysis using three variables, X, Y, and Z, which are all connected by pair-wise relationships (solid lines, left). Correlations describing specifically the relationships between X and Y, as well as X and Z (middle). The contribution of X to both Y and Z is then subtracted from each (grey arrows), and the relationship between the residual Y and Z is calculated, giving the partial correlation (red line, right). **(b-g)** Comparison between total and partial correlations for all combinations of Asef, Rho GTPase, and edge velocity (total correlation, black line; partial correlation, red line). Edge to GTPase correlations, controlled for RhoGEF input (upper). GTPase to RhoGEF correlations, controlled for edge input (middle). Edge to RhoGEF correlations, controlled for GTPase input (lower). (**h**) Role and spatio-temporal dynamics of Asef, Cdc42 and Rac1 in controlling the cell edge. Asef directly activates Cdc42 to control 30% of the effect of Cdc42 on edge motion. Asef also contributes indirectly to the activation of Rac1, but this contribution does not play a role in controlling the edge. Instead, other GEFs must be responsible for the motion-relevant signaling of Rac1. The relative timing and localization of each protein are shown. Note that activation of Asef is tightly localized and occurs earlier than Rac1 and Cdc42, supporting localized GTPase activation by Asef followed by GTPase diffusion or transport to the edge.

To quantify the relationship between biosensor activity and edge displacement, we used our previously described local cross-correlation analysis^3^, which integrates data from multiple cells to assess the significance of coupling between the two cues despite heterogeneous signaling and motion along the cell periphery. Specifically, we tracked the cell edge over time and then divided the perimeter of the cell into small edge sectors and associated sampling windows (Figure 2b). Sampling windows were placed in layers of increasing but constant distance from the edge. As the cell boundary moved, the windows were rearranged such that they maintained a stable one-to-one relationship with the associated sector. Thus, we could sample for each edge sector the local, instantaneous velocity and the corresponding biosensor activity at any layer (Figure 2c). These cell shape- and motion-invariant data representations then permitted a straightforward analysis of the relationship between edge motion and signaling activity using Pearson’s correlation (Figure 2d).

Improvements in instrumentation since our published studies enabled us to enhance spatial resolution by sampling 0.7 x 0.7 micron windows rather than the 1.4 x 1.4 micron windows used previously. Consistent with the earlier work in fibroblasts, Rac1 correlated best with edge motion in the two layers between 0.7-2.1 µm (Figure 2e, f). Also consistent with the earlier studies, correlation was strongest when we incorporated a lag between edge movement and Rac activity, but the lag seen in MDA-for MB-231 cells was considerably shorter than in fibroblasts (5s, Figure 2e, f). The shortest time lag to protrusion/retraction cycles was in these two middle layers, suggesting that Rac1 molecules are activated in this zone and then diffuse or are transported to the cell front. The correlation of Cdc42 with edge motion also peaked at a distance 1 – 2 µm from the cell edge, and with similar time lag, but with overall weaker significance than Rac1 (Figure 2e, f), suggesting that Rac1 is the more dominant GTPase in the regulation of MDA-MB-231 protrusion and retraction.

Asef had a maximum correlation in the first layer, immediately adjacent to the edge (0-1.4 µm, Figure 2e). Remarkably, and in contrast to the GTPases, we observed a near twofold increase in the correlation peak magnitude when we switched to analyzing correlation with the new, smaller windows, despite the decreased signal/noise that they produced (compare Figures 2e and 2f). This indicated that the portion of the Asef signal related to cell motion was activated in a highly confined zone 0.7-1.4 µm behind the cell edge, which corresponds to the zone of cell adhesion formation (**Extended Data Figure 4b**). Like Rac1 and Cdc42, the Asef activation was slightly delayed relative to protrusion-retraction cycles. Together, these results indicate that spatially precise RhoGEF activity produces more diffuse effector activation, probably due to diffusion of the effector from the point of activation. Unlike Asef, Vav2 showed no significant correlation with edge motion at the coarser analysis resolution (Figure 2e), and only a weak correlation in finer analyses using the 0.7-1.4 µm layer (Figure 2f). This indicates that Vav2 plays no major role in coordinating signals that specifically regulate cell protrusion and retraction events in unstimulated migration. Given the strong response of Vav2 to acute EGF stimulation (Figure 1h), we hypothesize that Vav2 is important in translating directional cues to the cell protrusion machinery.

In view of the significant correlation between Asef activity and cell edge movement we decided to focus on the interactions of Asef with the GTPases Rac1 and Cdc42. To complete our statistical analysis, we needed to correlate the activation of Asef and each GTPase, so we produced red shifted GTPase biosensors that could be imaged in the same cell as the Asef biosensor. We modified our published GTPase biosensors^3^ by changing the fluorescent proteins to LSSmOrange^34^ and mCherry^35^ (Figure 3a, b). Because of LSSmOrange’s long Stokes shift, we could capture emission from both biosensors at once using a single excitation, for rapid imaging and reduced photo-toxicity (Figure 3a, b). The new biosensors’ responded correctly to RhoGEFs and RhoGAPs, and their dynamic ranges were similar to those of the original blue-yellow biosensors (**Extended Data Figure 3**). The GTPase biosensors were transfected into cells stably expressing Asef biosensor, at levels that minimized perturbation of cell motility (**Extended Data Figure 6**). Cells with two biosensors showed activation patterns like those of single biosensor cells (by correlation analysis and by visual inspection, Figure 3b, **Extended Data Figure 7**). With two biosensors in the same cell we could employ the dynamic grid of sampling windows to extract the local correlation between Asef and each GTPase (Figure 3c, d). Asef activity correlated with both Rac1 and Cdc42, in both the 0-1.4 µm and 1.4-2.8 µm layers. Overall, both correlation peaks were significantly higher for Cdc42 than for Rac1. This is consistent with biochemical data that Asef directly interacts with Cdc42, but can activate Rac1 only indirectly ^9,36^, although our imaging data now shows that all three signals are active in the same zone. Any delay between activation of Asef and Cdc42 was too short to be discerned, given the 5s resolution necessitated by imaging both biosensors together, and the Asef-Cdc42 correlation was essentially symmetric about t=0. We noted that the overall positive correlation lobes in the second layer dip at t=0. Thorough investigation of the causes unveiled a systematic amplification of subpixel errors in cell edge segmentation and sampling window positioning that depresses the correlation values specifically at zero time lags **(Extended Data Figure 8).** Although the Asef-Rac1 and Asef-Cdc42 correlation peaks were close to t=0, there was a strong bias towards positive lag times (Cdc42 shows 55%, and Rac1 shows 67% after t=0). Therefore, relative to Asef activation, Cdc42 activation precedes Rac1 activation, which is again consistent with the hypothesis that Asef promotes Rac1 activation only indirectly

Taken together, this analysis revealed specific regions and times when activation of Asef is correlated with Cdc42 and Rac1 activation during cell edge movement. During protrusion, the GTPases Rac1 and Cdc42 are co-regulated by multiple RhoGEFs, which our data show include Asef but not Vav2. A critical question is why a RhoGEF activates multiple GTPases, and how much interaction with each one contributes to the downstream effector responses that actually produce edge motion. In concrete terms here, how much of the Cdc42 activation that is produced specifically by Asef contributes to the modulation of edge motion, and how much is contributed by Asef’s indirect activation of a Rac1 signal? By imaging all possible combinations of three variables (GEF/GTPase, GEF/edge, and GTPase/edge), we could address this question using partial correlation analysis.

Given three co-fluctuation variables (X,Y,Z), partial correlation analysis determines the level of direct coupling between fluctuations in Y and Z that are independent of the fluctuations driven by the coupling to the third variable X (Figure 4a). Using this approach, we first computed the correlation between Cdc42 activation and edge motion that is independent of Asef activity. We focused on the layer 1.4 – 2.8 µm from the edge, where Asef displayed significant co-fluctuation with edge motion and Cdc42 (Figures 2e, 3d). Compared to the direct correlation between Cdc42 and edge motion, the peak value was about a third smaller (0.2 versus 0.13) (Figure 4b), indicating that about 33% of the Cdc42 signal that influences edge motion is triggered by Asef. In stark contrast, for Rac1 the correlation with edge motion increases after eliminating the contribution of Asef (Figure 4c). This means that the transduction of signal from Asef to Rac1 dampens the co-fluctuation between Rac1 and edge motion.

This surprising finding is again consistent with indirect activation of Rac1 by Asef via cross-talk between Cdc42 and Rac1. Alternatively, interactions between Asef and Rac1 mediated without Cdc42 may be elicited for cell functions other than protrusion, again with an adversarial effect on the interactions between Rac1 and edge motion. Distinguishing these two explanations will require concurrent imaging of Asef, Cdc42, and Rac1, which is impossible at this point. Nonetheless, these findings reveal the limitations of multi-functional signaling networks, where signaling for one purpose may impair the precision in signaling for another purpose. Previous studies suggest that expansion of the plasma membrane during protrusion produces mechanical feedback on GEF-GTPase interactions, especially in the adhesion-rich zone near the cell edge^37,38^. We therefore applied partial correlation analysis to assess the influence of cell edge motion on Asef/Cdc42 and Asef/Rac1 relationships. Both these interactions were nearly independent of edge motion (Figure 4d, e), indicating that Asef is unlikely to be a mediator of mechanical feedback to Rac1 and Cdc42.

Finally, we asked how eliminating the influence of the GTPases would affect the correlation between Asef and edge motion. For both Cdc42 (Figure 4f) and Rac1 (Figure 4g) the correlation curves substantially decreased, confirming that the interaction between Asef and motion is indeed critically mediated by Rac1 and Cdc42. Intriguingly, with Cdc42 effects removed, the Asef/motion correlation was markedly reduced at negative time lags and between 0 and 25 s (Figure 4f), whereas removal of Rac1 inputs reduced the Asef/motion correlation at lags between 0 and 50s (Figure 4g). This suggests that coupling of Asef to edge motion early in protrusion is dependent on Cdc42, while later coupling depends more on Rac1. Again, this finding is consistent with our previous interpretation that Asef activates Cdc42 directly, but Rac1 is coupled to Asef via an indirect connection, leading to a delay of ~25s.

In summary, the study of RhoGEFs has been limited to deciphering biochemical interactions *in vitro*, and to molecular perturbation *in vivo*^39,40^. However, due to the complexity of RhoGEF-GTPase interaction networks, which contain numerous feedbacks, crosstalks and redundancies, an understanding of GEF function requires new methods to determine connectivity and function in the context of space and time. To unravel the effects of RhoGEFs on multiple GTPase targets, we first developed a generalizable approach to Dbl family GEF biosensors. The method is applicable to diverse structures comprising 19 of the 70 Dbl family RhoGEFs (**Extended Data Figure 2d**), and potentially to 10 GEFs that are similarly regulated. This enabled correlation of GEF activity and GTPase activity with the motions of the cell edge. We then modified our existing GTPase biosensors for simultaneous imaging of GEF and GTPase activity, providing the ability to directly correlate their spatio-temporal dynamics in living cells. With pairwise correlation of edge motion, GEF activity and GTPase activity in hand, we could deploy partial correlation analysis to locally sampled time series and determine that the RhoGEF Asef has stronger interactions with Cdc42 than with Rac1, albeit both interactions are statistically significant. We could show that specific, spatially localized Asef activations contribute to specific Cdc42 activation events, and determine the percentage of Cdc42 activation relevant to edge activity that stemmed from Asef (Figure 4h). For Asef activation of Rac1, several observations (timing, diffuse response, and weaker coupling between Asef and Rac1) indicated that Rac1 is activated via indirect interactions, consistent with published biochemical studies. Moreover, while Asef activity clearly has an activating effect on Rac1 signaling, our partial correlation analysis indicates that the Asef-triggered Rac1 modulation does not contribute to the control of edge protrusion. This highlights that each signal had to be understood through *in situ* analysis in the context of a cell function. Building on these technologies, future work will now allow the comprehensive functional assignment of the perplexing multitude of RhoGEF-GTPase interactions in the context of diverse cellular behaviors.

## Methods

### RhoGEF biosensor design

The biosensors were optimized and tested in a sequential fashion. Initially, a cassette comprising mCerulean or mCerulean3, a flexible linker^7^, and mVenus or YPet was inserted into a series of sites between the AID and DH domains of full-length RhoGEFs and emission was measured using fluorometry for Vav2 or high-throughput microscopy for the other RhoGEFs (see detailed procedures below). For Vav2 we compared wild type to constitutively active (Y140E:Y159E:Y172E, 3YE) mutants, and for the other RhoGEFs we used co-expression of an activator. For Vav2 we also tested linker variants. These comprised a flexible unit^41^ and a structured helical unit^42^ combined in different topologies and repeat numbers (2, 3, and 4 repeats for short, medium and long respectively). Once we had identified the best insertion site and linker combination we combinatorially combined donor proteins (mCerulean3, TagCFP and mTFP) with acceptor proteins (YPet, along with circular permutations of YPet) to form a library of variants of each biosensor that were screened using high-throughput microscopy for the effects of activation. The biosensor constructs were inserted into a tet-off inducible retroviral expression system and stable lines were produced in tet-off MDA-MB-231 cells (ATCC). Cells were maintained in DMEM (Cellgro) with 10% FBS (Hyclone) and 0.2 μg/ml doxycycline to repress biosensor expression. Biosensors were named FLARE.a as part of a nomenclature system based on the biosensor design (eg Asef FLARE.a)

### Rho GTPase biosensors

The Rac1 FLARE.dc1g biosensor is a modification of our previously reported dual chain biosensor design^3^. To improve brightness and dynamic range we used Turquoise fluorescent protein^13^ rather than CyPet. In brief, YPet^43^ is fused upstream of residues 60 −145 of human PAK1, and Turquoise is fused to the N-terminus of full-length Rac1. The two biosensor chains were expressed on one open reading frame with two consecutive 2A viral peptide sequences from *Porcine teschovirus*-1 (P2A) and *Thosea asigna* virus (T2A) inserted between them, leading to expression of the two separate biosensor chains^44^. The Cdc42 and RhoA FLARE.dc1g biosensors use an identical topology, wherein YPet is fused upstream of N-WASP (200-293) or Rhotekin (6-98), for Cdc42 and RhoA respectively, and mCerulean3 is fused to the N-terminus of the GTPase, with the two biosensor chains separated by the 2A sequences. The biosensor constructs were inserted into a tet-off inducible retroviral expression system and stable lines were produced in tet-off MDA-MB-231 cells (ATCC). Cells were maintained in DMEM (Cellgro) with 10% FBS (Hyclone) and 0.2 μg/ml doxycycline to repress biosensor expression

For the red shifted versions of the GTPase biosensors, YPet was exchanged for mCherry and the donor was exchanged for LSSmOrange. Both donor and acceptor contain the R125I mutation to increase intramolecular FRET^45^. For dual biosensor experiments, these red-shifted GTPase biosensors were transfected into the RhoGEF biosensor-expressing stable cell lines.

### Spectral analysis of biosensors

Emission spectra of biosensors were obtained using a Fluorolog fluorometer (Horiba). HEK-293t cells grown in 6-well plates (Nunc) were transfected with biosensor DNA plus regulator if required. After 24 h cells were detached by trypsinization (Cellgro) and resuspended in cold PBS (Sigma) + 1%FBS (Hyclone), washed and then resuspended in cold PBS. Samples were excited at 430nm and spectra obtained from 460 to 600nm for biosensors with Cerulean3/TagCFP/mTFP, and 550 to 650nm for LSSOrange biosensors. Dual chain biosensors were corrected for acceptor bleedthrough.

### High-throughput microscopy screening

Using a modification of our published procedure^10^, HEK-293t cells were plated in 96-well plates with µ-clear plastic bottoms (Greiner bio-one) coated with poly-l-lysine (Sigma). Cells were transfected in triplicate with a library of biosensor DNA plus regulator if required and imaged after 24 h. Growth media was replaced with HBSS (Sigma) with 1%FBS and 10mM HEPES (Gibco) prior to imaging. Cells were imaged using a 10X, 0.4 NA objective on an Olympus IX-81 inverted microscope and using Metamorph screen acquisition software (Molecular Devices) and mercury arc lamp illumination. Filters used were Ex - ET436/20X, Em; donor- ET470/24M, FRET - ET535/30M and a 445/505/580 ET dichroic mirror. Images were obtained on a Flash4 sCMOS camera (Hamamatsu). Images were analyzed using MATLAB (Mathworks). Briefly, 4 fields were taken of each well and the intensity was summed for each channel. These were then background subtracted using values from wells that were mock transfected, and ratios obtained from these background subtracted values. For regulator titration experiments, regulator DNA was first titrated against the biosensor DNA and then transfection complexes formed in 96-well plates prior to transfection. Regulators used: APC (309-798), Src (FL, Y529F), Gα_13_ (FL, Q226L), PAK (FL D389R:S422D:T423E), Dbl (495-826), Vav2 (191-573), Asef (FL), Tiam1 (C1199), p115RhoGEF (FL), RhoGDI (FL), p50RhoGAP (FL), RacGAP1 (FL).

### Src stimulation of Vav2 biosensors

Cells were transfected and imaged as above. After lysis, triplicate samples were pooled and Western blotted for phospho-Vav levels using a phospho-specific Tyr-172 antibody (Abcam). Samples were normalized to biosensor expression using an anti-GFP antibody (Clontech).

### Stimulation experiments

A431 cells (ATCC) were plated in 6-well plates 24hr prior to transfection. Cells were incubated with transfection complexes for 5-6 hours and then replated onto #1.5 coverslips (Warner Instruments) coated with Collagen IV (Gibco). Cells were imaged in DMEM/10% FBS. After 2-3 h the media was replaced with DMEM/0.5% delipidated BSA and cells were starved overnight prior to imaging. Cells were then imaged in Hams/F12 (Caisson Labs) with 0.5%BSA, 10mM HEPES (Gibco), 100 µm Trollox (Sigma), and 0.5mM Ascorbate (Sigma). Cells were imaged using a 40x 1.3NA Silicon oil objective on an Olympus IX-81 inverted microscope using Metamorph software and 100 W Hg arc lamp illumination. Excitation filters used were FF-434/17 and FF-510/10 combined with a FF462/523 dichroic mirror. Donor and FRET images were simultaneously captured using a TuCam system (Andor) fitted with FF-482/35 and FF-550/49 and an imaging flat FF509-FDi01 dichroic, together with two Flash4 sCMOS cameras (Hamamatsu).

### Constitutive migration experiments

Biosensor expression was induced 48 hr prior to imaging through trypsinization and culturing without doxycycline. On the day of imaging, cells were replated using Accumax (Innovative Cell Technologies) onto coverslips coated with collagen I (10µg/ml 37C overnight) and allowed to attach in DMEM /10%FBS. After 2hrs the media was replaced with Hams/F12 with 0.2% BSA, 10ng/µl EGF (R and D systems) 10mM HEPES, 100 µm Trollox, and 0.5mM Ascorbate and cells were allowed to equilibrate. After a further 2-4 hrs, cells were imaged in a closed chamber with media treated with Oxyfluor (1/100). For single biosensor experiments, cells were imaged using the filters listed above. For dual biosensor experiments, the excitation filters used were FF-434/17 for Cerulean3/mTFP and LSSOrange, and FF-546/6 for Cherry (Semrock) combined with a custom zt440/545 dichroic (Chroma). For emission, a TuCam was fitted with a FF560-FDi01 imaging flat dichroic and a Gemini dual view (Hamamatsu) was added to each emission port. For the short wavelength Gemini the filters used were donor- FF-482/35, FRET - FF-520/15 and a FF509-FDi01 imaging flat dichroic mirror. For the Red-shifted Gemini the filters used were Orange – FF01-575/15, FRET/mCherry – FF01-647/57 and a FF580-FDi01 imaging flat dichroic.

### Image Processing and Analysis

Biosensor activation levels were measured in living cells by monitoring the ratio of FRET to donor emission on a pixel by pixel basis (6). Donor and FRET images were aligned using fluorescent beads as fiduciaries to produce a transformation matrix using the Matlab function “cp2tform” (Matlab, The Mathworks Inc.). This was then applied to the Donor image using the Matlab function “imtransform”. The camera dark current was determined by obtaining images for each camera without excitation, and the dark current was subtracted from all images. Images were corrected for shading due to uneven illumination by taking images of a uniform dye solution under conditions used for each wavelength, normalizing this image to an average intensity of 1 to produce a reference image for each wavelength, and then dividing the images corrected for dark current by the shading correction reference image. Background fluorescence was removed by subtracting, at each frame, the intensity of a region containing no cells or debris. Images were segmented into binary masks separating cell and non-cell regions using the segmentation package “MovThresh” (7), which is based on the Otsu algorithm (8). The Donor channel was used for segmentation, as it had the highest signal to noise, particularly at the cell edge. The masks were then applied to all channels, setting non-cell regions to zero intensity. For dual biosensor imaging these masked images were then corrected for bleedthough of Cerulean3/mTFP and YPet into the orange/red channels.

For RhoGEF biosensors, activation maps were obtained by dividing the corrected donor image by the FRET image. For the GTPase biosensors, the images were corrected for bleed-through and ratios were obtained using the following equation (using data from control cells expressing donor or acceptor alone to obtain the bleed-through coefficients α and β): R= (FRET – α(Donor) – β(Acceptor))/donor where R is the Ratio, FRET is the total FRET intensity as measured, α is the bleed-through of the donor into the FRET signal, β is the bleed-through of acceptor into the FRET signal, and Donor and Acceptor are the donor and acceptor intensities as measured through direct excitation. These ratio images were then corrected for photobleaching. For stimulation experiments, the ratios were divided by a reference curve derived from mock-stimulation experiments. For constitutive migration the whole cell average was fitted to a double exponential curve and this curve was used to normalise. Pseudocolor scales were produced without considering the lowest and highest 5% of ratio values to eliminate spurious pixels, and normalizing so the lowest value is 1.

### Cell windowing analysis

After the corrections described above, the cell images of the ratiometric biosensor activity were compartmentalized into layers of rectangular windows along the entire cell edge. To construct the sampling windows at a constant distance from the cell edge we computed a distance map (Matlab function: bwdist) to the segmented cell edge. The distance map yielded equidistant contours at either 0.7µm (2 pixels) or 1.4µm (4 pixels) from the cell edge. At the cell edge the first contour was divided into segments of 1.8 - 3 μm width (Figure 2b). Sampling windows located at the cell edge derived a time course of edge velocity and biosensor intensity as the windows tracked the morphological changes of the cell over time. For windows placed in layer 2 and higher only the biosensor intensities were sampled. However, each of these windows maintained unambiguous correspondence to a window at the cell edge, allowing correlation of biosensor intensity fluctuations inside the cell with cell edge movements. Importantly, the ability to maintain unique correspondences depended on the cell morphology. Windows for which the correspondence was lost at one or several time points of a movie because of particularly strong cell morphological changes were eliminated from the analysis.

This *in silico* compartmentalization of the cell allowed us to represent the biosensor activity into a cell-shape invariant space. For each frame of the movies, the biosensor activity was averaged within the area of each sampling window, resulting in a set of matrices representing the biosensor activity of a layer of windows with a fixed distance from the cell edge. Rows correspond to windows and columns to time (Figure 2c). This method has shown to be an efficient way to spatiotemporally sample the activity of sensors expressed by the cell. For more information, see^46^.

### Windows selection

Migrating cells usually display regions along the cell edge that are active with protrusion/retraction cycles, but they also exhibit quiescent regions with little morphodynamic activity. We implemented window selection criteria based on the autocorrelation function of the protrusion/retraction speed estimated by the windowing algorithm described above. The autocorrelation function of a random variable *X* can be described as the cross-correlation between *X* and its time-delayed version *X (lag)* where *lag* is the duration of the delay. It can measure an average duration of “memory” in the signal that can be described as the maximum time spacing between samples that still exhibit a linear association. No significant linear correlation can be measured when taking samples further than this duration apart. Consequently, the autocorrelation function of a signal with structure has a much slower decay compared with the autocorrelation function of a signal with samples independently drawn from a uniform random distribution. We used the full width at half maximum (fwhm) of the window speed autocorrelation as a measure of information in the signal that our analysis methods can make use of. Only windows with autocorrelation fwhm equal to or higher than 0.6 frames were chosen for analysis, i.e. the speed time series in such windows showed on average some temporal coherence in motion. For reference, a white noise time series with no coherence has a fwhm < 0.5 frames. The effect of this strategy was the elimination of quiescent regions. For the example in Figure 2, all windows are shown in **Extended Data Figure 9**.

A second method of window selection was implemented in order to avoid a phenomenon we observed when cells were expressing two biosensors. This problem arose due to a sub-pixel segmentation error that appears when two biosensors have opposing gradients. See **Extended Data Figure 8**. This transient error causes a strong negative cross-correlation between the activities of the two imaged biosensors. In order to minimize the impact of this artifact on our final analysis, we excluded windows that presented a sharp negative cross-correlation between two biosensors at lag zero. It is worth mentioning that windows with a negative cross-correlation trend that resulted in a negative score at lag zero were not excluded from analysis. The selection algorithm starts by decomposing the cross-correlation between the biosensors for a given window using empirical mode decomposition - EMD^47^. This decomposition technique recursively extracts components of the signal from the fastest, or higher frequency content, to slower component of the signal. The fastest component or first intrinsic mode function (IMF) absolves all fast variations present in the signal. The window is excluded from analysis if after its cross-correlation decomposition, the amplitude of the first IMF at lag zero exceeds a threshold and shows change in derivate for the lag zero neighborhood. The threshold is estimated as three standard deviations away from the mean using the first IMF points to build the distribution.

### Pearson correlation analysis

The main aim of the Pearson correlation analysis is to determine the strength of the linear relationship between two random variables. For instance, it can be applied to find time-dependent linear relationship of two measured parameters, which is the case in this work. The correlation value ranges from [-1,1] where score value 1 represents perfect linear correlation, 0 represents no correlation and −1 perfect anti-correlation. Additionally, one time series can be time-shifted in relation to the second by a value usually referred to as *lag*. This lag analysis can be useful when two variables are correlated with a time delay. That means there will a value of lag where the correlation score is maximum. The lag at peak correlation can then be interpreted as the time needed for one variable to process upstream information or as the time taken for information to be transferred or a combination of both. Because correlation analysis requires a stationary signal, few pre-processing steps are required before the correlation coefficient is calculated. All time series data used to calculate a correlation score in this work were mean subtracted and linear trend removed.

Pearson’s correlation coefficient *ρ*(*a*_1_(*t*), *a*_2_(*t*))_*τ*_ between two activity time courses *a*_1_(*t*) and *a*_2_(*t*) was computed as a function of the time lag *τ* using the Matlab function (xcov). This function implements the mean corrected and normalized correlation functions as follows:

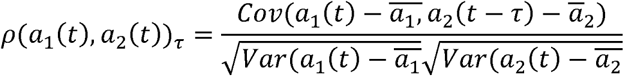

The operators Cov(.) and Var(.) denote the covariance and the variance of the mean corrected time courses, respectively. The variables 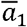 and 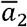 denote the mean value of the respective time course. Correlation functions were first calculated for pairs of time courses per window (either between edge velocity and biosensor intensity, or between two biosensor intensities in the case of simultaneous imaging GEF and GTPase activity). For each correlation curve we computed the confidence level at which correlation became significant relative to the expected correlation of random time series. This level (~0.1 for all our data) depended on the duration of the movies and the number of sampling windows. In addition, through bootstrap analysis of the variation among all correlation curves, we computed confidence intervals for the mean cross-correlation. This reflected the number and consistency of sampling windows across experiments. This computation is performed by Fisher transforming the correlation values with a hyperbolic tangent and bringing back the distribution parameters with the hyperbolic tangent inverse^48^.

### Partial correlation analysis

Although Pearson correlation analysis has been shown to be useful when only two variables are at hand, it can result in misleading outcomes when trying to identify linear relationships between random variables where the data generating system has confounding structure^49^. A simple example of this case is when a random variable *A* drives two other unrelated random variables *B* and *C*. There is a flow of information going from *A* to *B* and *A* to *C* but no exchange of information between *B* and *C*. A simple correlation analysis would identify the confounding information from *A* shared by *B* and *C* as a direct connection between *B* and *C*. However, the partial correlation between *B* and *C* control for *A*, or *partial(B,C|A)*, would statistically return zero. For this simple three variable example, the partial correlation can be computed as:

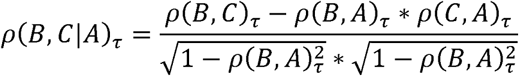

The time dependency of the random variables was omitted for simplicity. The equation above shows that partial correlation removes the influence of the variable A from B and C and renormalizes it such the final value lies within the [-1,1] interval. Although simple, using the inverse of the correlation matrix is a more efficient way to calculate the partial correlation when more than 3 variables are present^49^.

Because of the stationary requirement, all time series were mean subtracted and linear trend corrected as was done for the simple correlation analysis. Confidence intervals and levels were calculated in the same fashion as the Pearson’s correlation.

---

**Extended Data Figure 1**

**(a)** Screening RhoGEF biosensor designs. For each RhoGEF: Upper - Screening donor (wedge color) and acceptor (Wt – YPet; CP1 – YPet CP157; CP2 - YPet CP173; CP3 – YPet CP229) fluorophore combinations. Lower - Screening different insertion sites for the fluorescent protein cassette. **(b)** Comparison of different donor to acceptor linker compositions for the Vav2 biosensor. **(c)** Emission spectra of the variant with the highest dynamic range for each donor. Donor/FRET emission ratio change between active and wild type conformations is indicated, ex = 430nm. **(d)** Emission spectra of biosensors for Vav1 and Vav3 (∆R/R_off_ = Donor/FRET emission ratio change between active and wild type mutants, ex = 430nm).

**Extended Data Figure 2**

**(a)** A431 cells expressing Tiam1 biosensor after stimulation with EGF (100ng/µl). Donor/FRET ratio images pseudo-colored with scale bar at bottom right. **(b)** Whole cell average Donor/FRET ratio of A431 cells expressing the Vav2 biosensor before and after stimulation with the indicated concentrations of EGF. Graphs are average of 10 cells per concentration, error bars indicate 95% confidence intervals. **(c)** LARG (left) and β-Pix (right) activation reported by biosensors in MDA-MB-231 cells undergoing constitutive edge motion. Arrows point to activity in protrusions. **(d)** DH family RhoGEFs amenable to biosensor production through modification of the hinge region

**Extended Data Figure 3**

Response of GTPase biosensors to upstream regulators. Biosensors and regulators were co-expressed in HEK293T cells and analyzed using high-throughput microscopy. Green bars represent positive regulators, red bars negative regulators and yellow bars are RhoGEFs that should not directly activate. Bars represent mean of 6-9 independent transfections across multiple experiments. Error bars are 95% C.I.

**Extended Data Figure 4**

**(a)** Relationship between biosensor expression level and average protrusion velocity. **(b)** Relationship between biosensor expression level and peak correlation of biosensor activity with edge velocity. No significant relationship was detected between biosensor expression level and velocity or correlation at these expression levels. Expression levels were determined relative to the fluorescence of the medium in the shade-corrected FRET image. Grey box indicates expression levels where the SNR is too low for FRET analysis.

**Extended Data Figure 5**

**(a, c, e)** Asef activation reported by biosensors in MDA-MB-231 cells undergoing constitutive edge motion (right). Cells co-express markers labelling **(a)** F-Actin (F-tractin), **(b)** Focal adhesions (Paxillin), and **(c)** Late endosomes/sorting components (Rab7) (left). Arrowheads point to activation at the edge, Arrows point to second layer for Asef that lies behind lamella/adhesion zone. **(d)** Control biosensor for Asef shows greatly reduced ratio across the cell.

**Extended Data Figure 6**

**(a)** Comparison of GTPase biosensor expression using Cerulean3/YPet and LSSOrange/mCherry based biosensors. Relative expression was calculated using donor brightness, adjusting emission intensity for differences in dichroic, emission filter, protein brightness and camera efficiency at the different wavelengths (data obtained from fpbase.org). **(b)** Relationship between biosensor expression level and average protrusion velocity. Expression levels of both biosensors in each cell plotted. No significant relationship was detected between biosensor expression level and velocity or correlation at these expression levels.

**Extended Data Figure 7**

Average cross-correlation functions for each combination of edge, RhoGEF activity, and Rho GTPase activity for each biosensor pair (**(a-c)** Asef and Cdc42, **(d-f)** Asef and Rac1); (n = 6 Asef/Cdc42 cells, n = 6 Asef/Rac1 cells). Analysis using 0.7μm window size, 1.4-2.8µm layer.

**Extended Data Figure 8**

Diagram to illustrate potential artefacts caused by sub-pixel errors in segmentation of biosensor images when two biosensors have inverse gradients.

Thorough investigation unveiled an unfortunate systematic amplification of subpixel errors in the cell edge segmentation and window positioning that yields a depression of the correlation values specifically at zero time lags. When edge is correctly assigned (middle) the biosensor activity is correctly measured and the cross correlation is correctly measured. When the first window edge extends onto non-cell areas (top), both activities are low in the first window, so giving an incorrect positive correlation. In layer 2 and deeper, biosensor A (red) is artificially high, while biosensor B (blue) is low, giving a negative correlation at lag=0. When the first window edge fails to extend to the cell edge (lower), biosensor A (red) is artificially low, while biosensor B (blue) is high in all layers, giving a negative correlation at lag=0. The effect is strongest in the second window layer because of the inverse spatial gradients in RhoGEF and GTPase activity (RhoGEF activity increases with distance from the edge; GTPase activity decreases with distance from the edge). At this point, there is no remedy for this effect, though improvements in resolution and segmentation could ameliorate it.

**Extended Data Figure 9**

Comparison with Figure 2, showing excluded quiescent windows. **(a)** Map of edge velocity along the edge. Green regions are protruding, purple regions retracting. **(b)** Map of Asef activity along the edge. Red/yellow regions have high activity, blue regions have low activity. **(c)** Cross correlation coefficients between edge velocity and Asef activity. Gold shows high correlation, blue negative correlation. **(a-c)** Each column is a single time point along the edge, each row is a single position through time. Black boxes show quiescent regions that are excluded.

**Supplemental Movie 1**

Cdc42 (left) and Rac1 (right) activation reported by biosensors in MDA-MB-231 cells undergoing random edge motion. Pseudocolor as in Fig. 1.

**Supplemental Movie 2**

Asef (left, middle) and Vav2 (right) activation reported by biosensors in MDA-MB-231 cells undergoing random edge motion. Enlarged view on left shows activation of Asef at cell edge. Pseudocolor as in Fig. 1.

**Supplemental Movie 3**

Simultaneously imaged RhoGEF and Rho GTPase biosensors in MDA-MB-231 cells undergoing constitutive edge motion. Upper images show activation of Asef (left) and Cdc42 (right), lower images show Asef (left) and Rac1 (right). Pseudocolor as in Fig. 1.

## Acknowledgements

National Institutes of Health (NIH) Program Project Grant P01 GM103723 (KMH, JS, and GD), R01-GM062299 (JS), R01 GM071868 (GD), and GM-R35GM122596 (KMH) supported this work. DM was supported by funding from the Leukemia and Lymphoma Society (LLS). GG was supported by the “Integrated Training in Cancer Model Systems” Training Program, T32CA009156.

## Author contributions

DM, JS and KH designed the biosensors. DM carried out all experiments, except for work by JR in building Tiam1, Tim, B-Pix and LARG biosensors. GG and MA contributed to initial design of Tim and LARG biosensors respectively. MV and GD developed and carried out computational analysis. GD, JS and KH provided intellectual input in all phases of the study and directed the work. The paper was written by DM, MV, GD, JS and KH, with input from all authors.

## Author Information

Correspondence and requests for materials should be addressed to, or

