## Supplementary figures and images for "Connectivity analysis of GEF/GTPase networks in living cells"

### Supplemental Figures

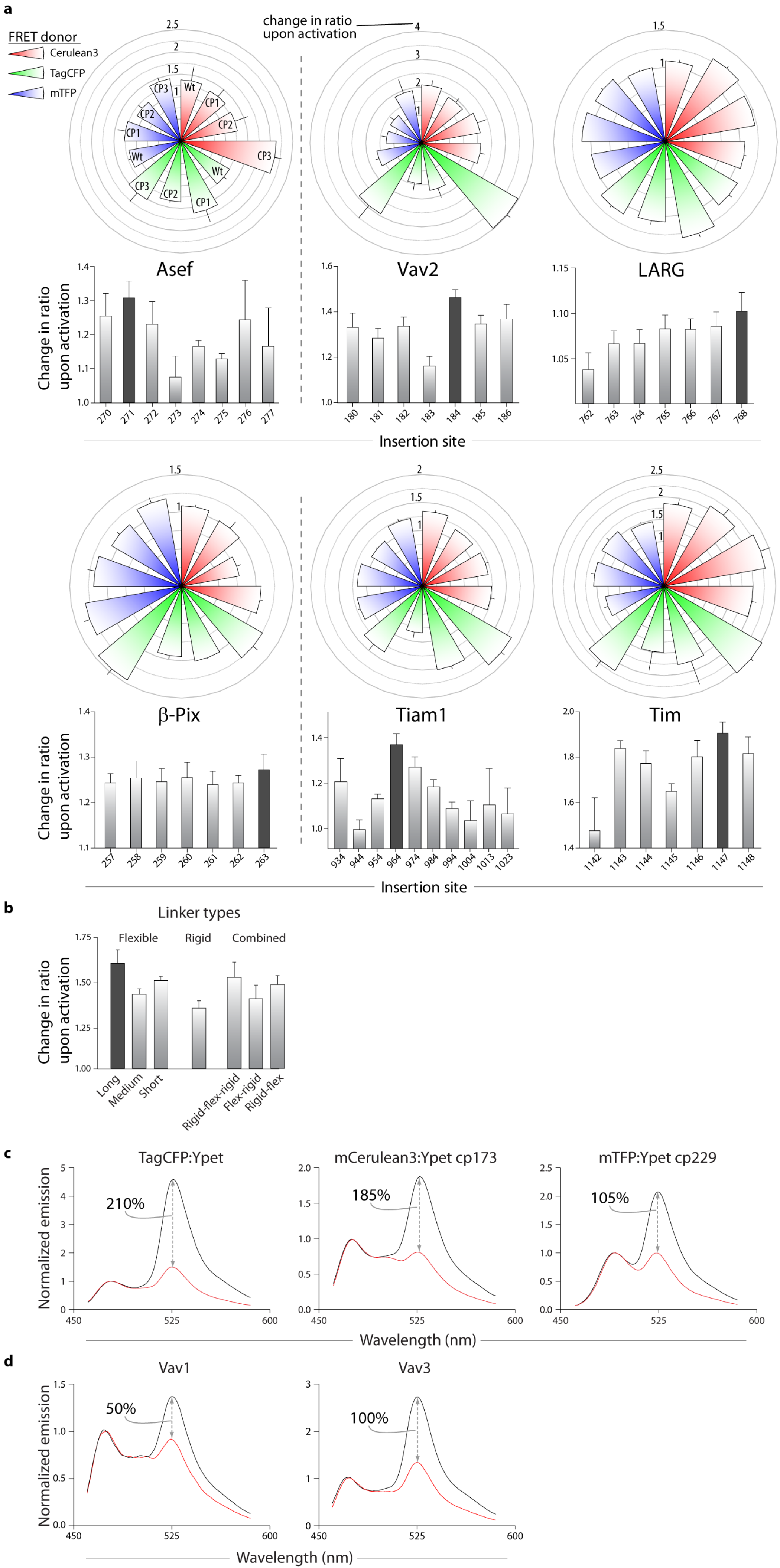

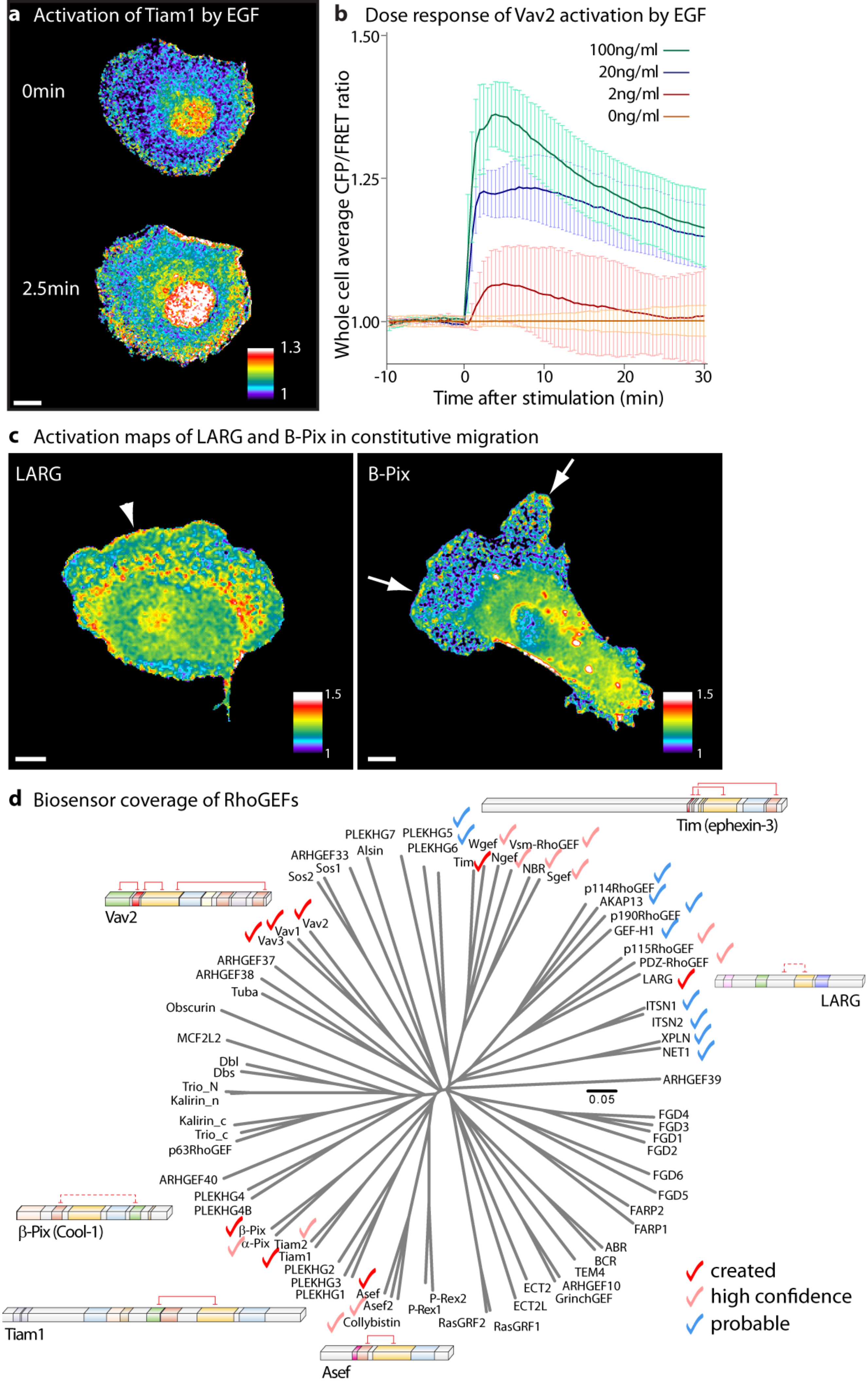

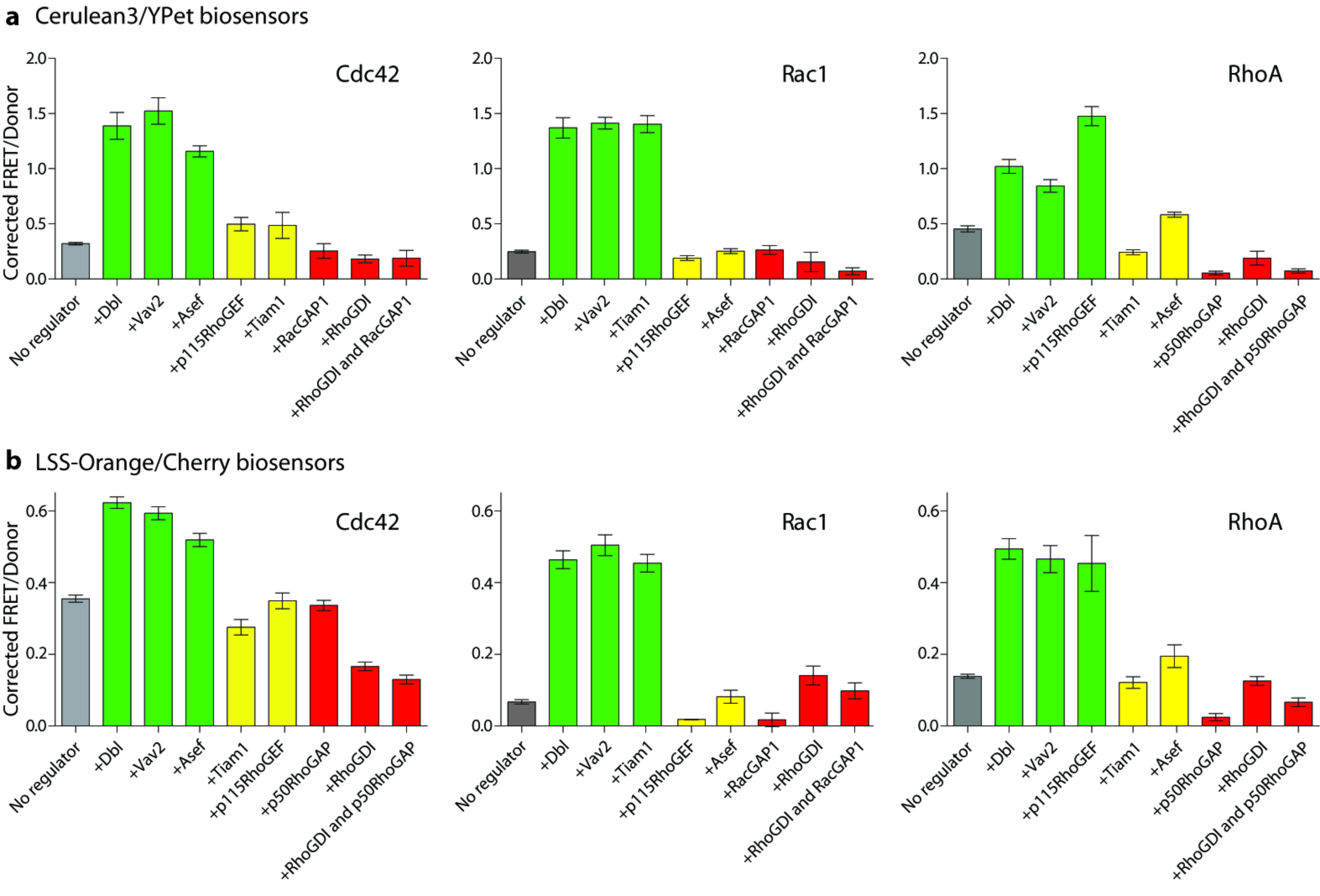

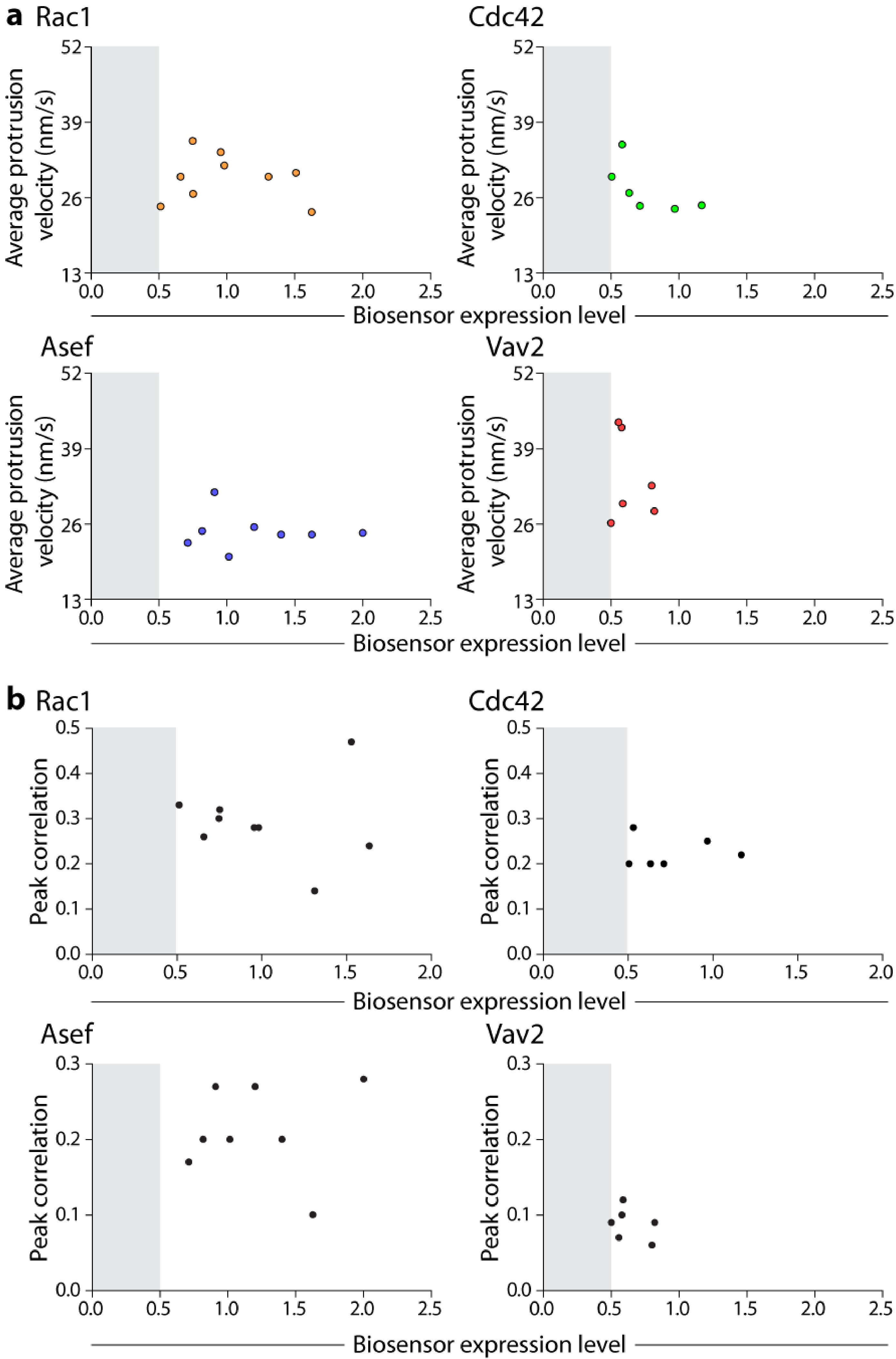

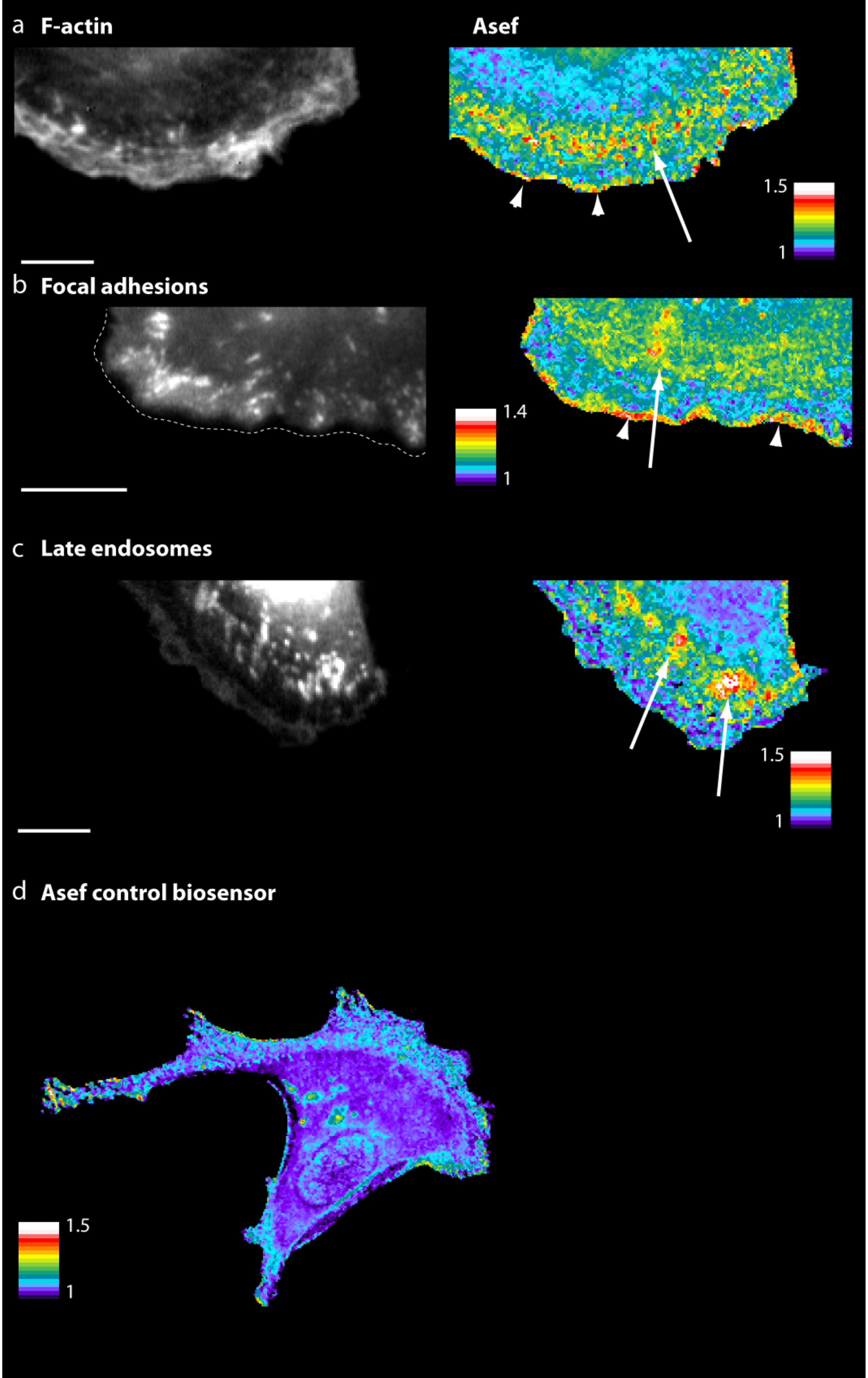

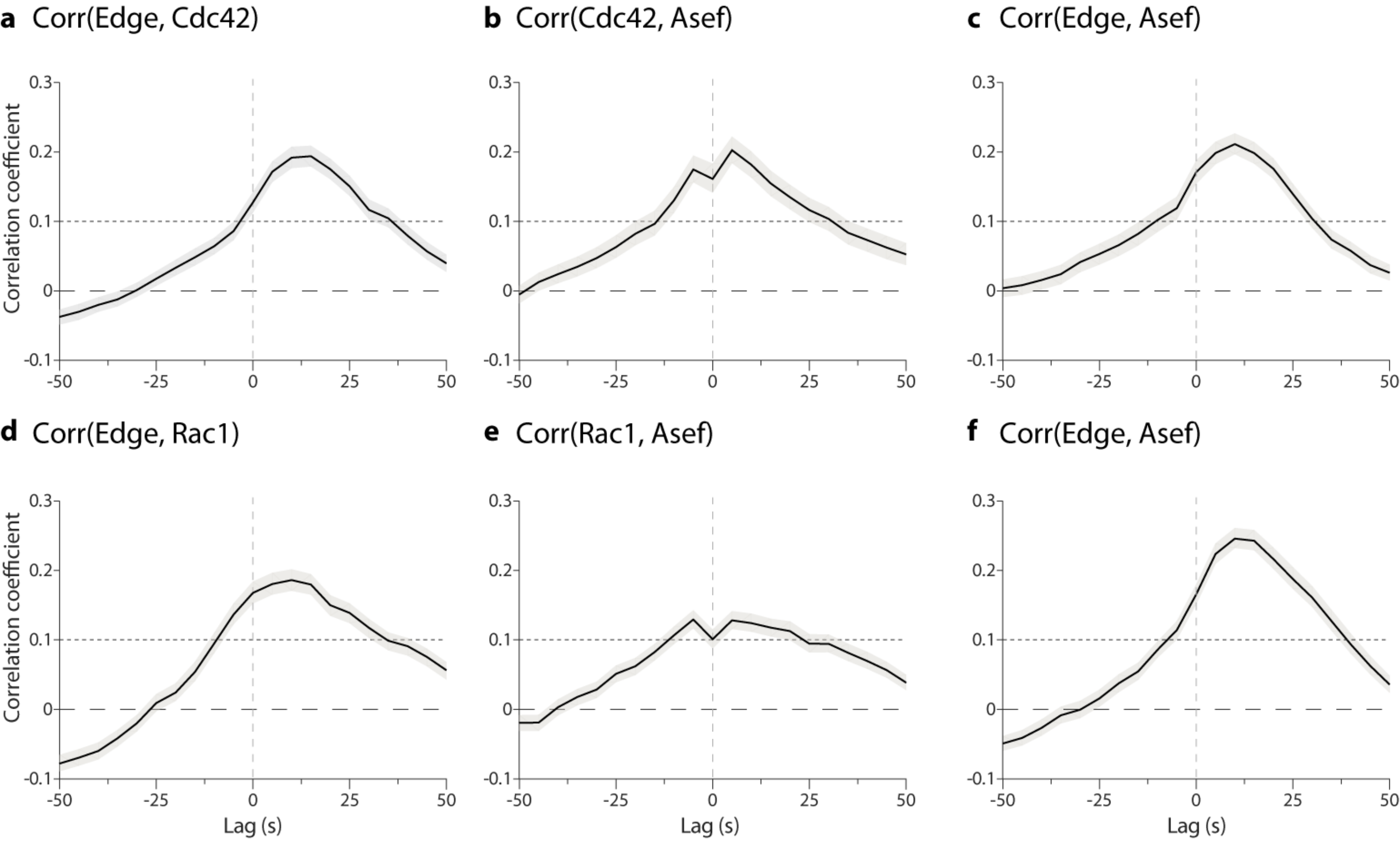

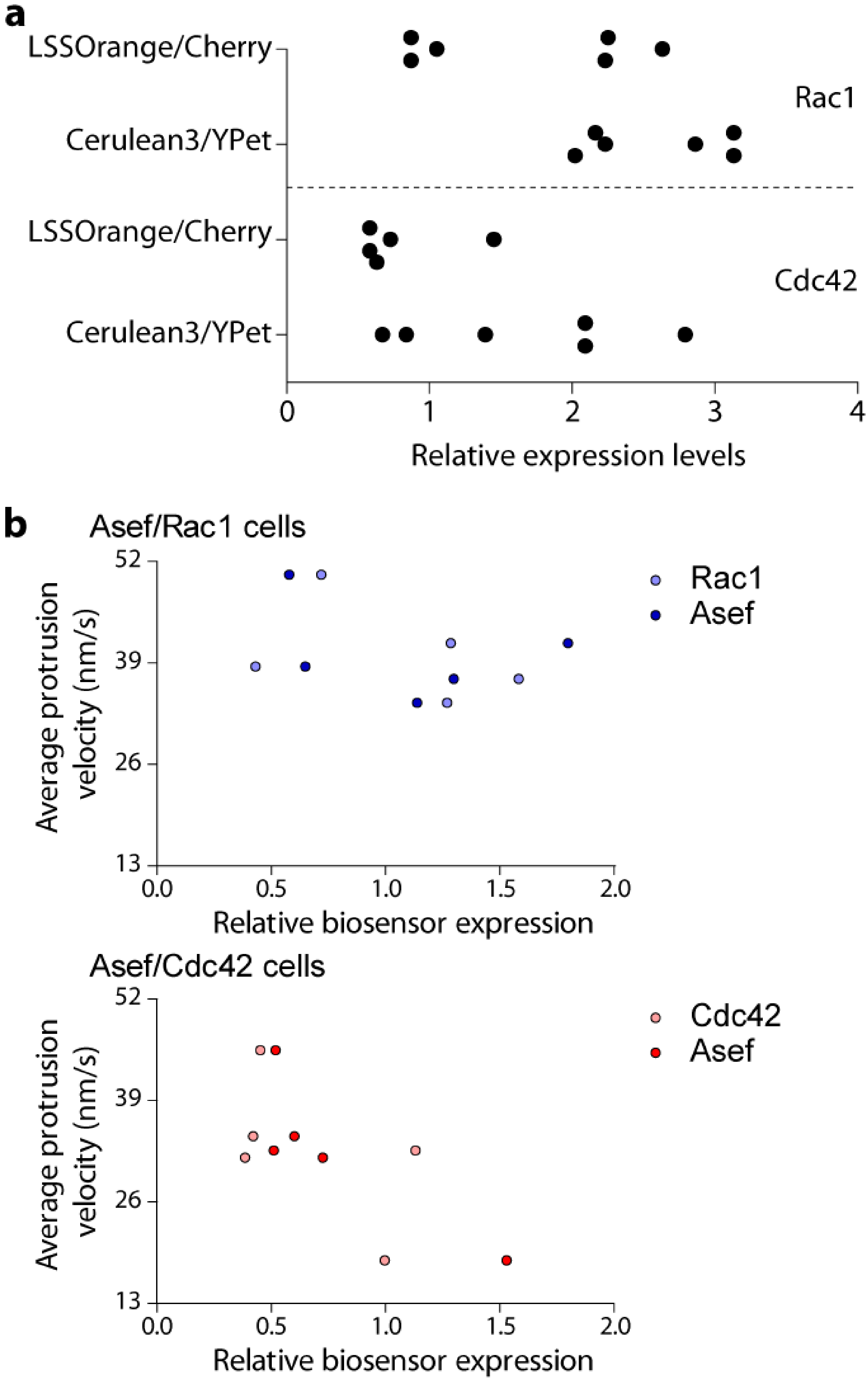

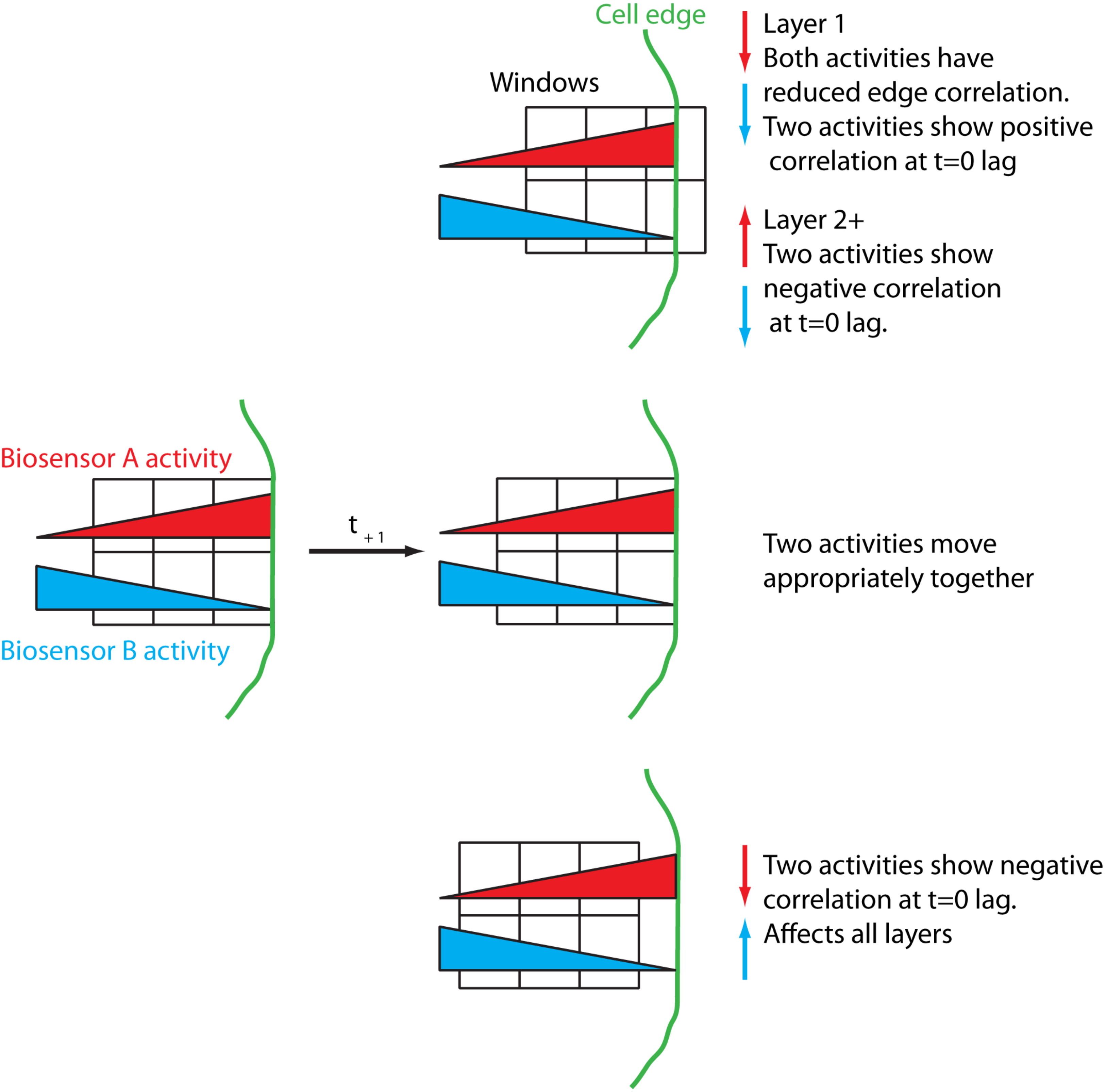

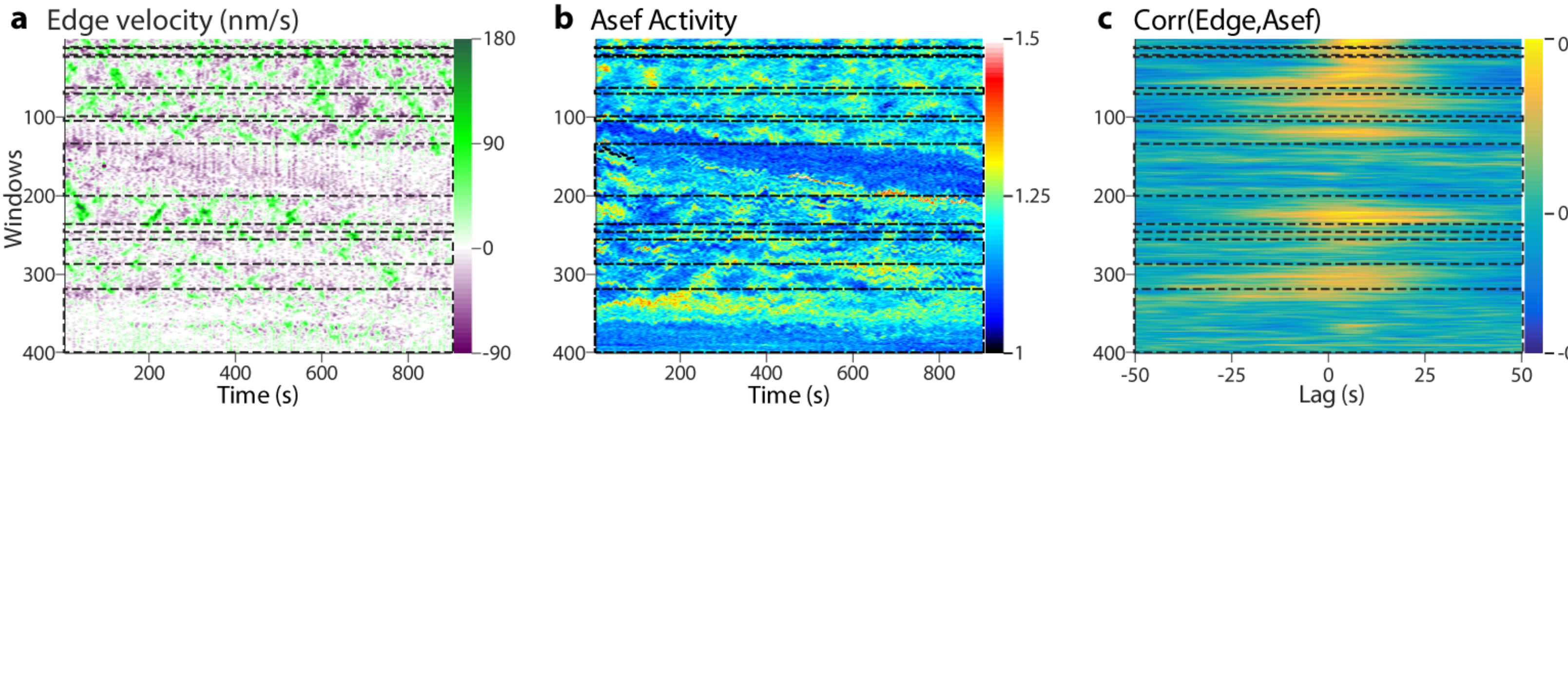
